# The Observed T cell receptor Space database enables paired-chain repertoire mining, coherence analysis and language modelling

**DOI:** 10.1101/2024.05.20.594960

**Authors:** Matthew I. J. Raybould, Alexander Greenshields-Watson, Parth Agarwal, Broncio Aguilar-Sanjuan, Tobias H. Olsen, Oliver M. Turnbull, Nele P. Quast, Charlotte M. Deane

**Affiliations:** Oxford Protein Informatics Group, Department of Statistics, University of Oxford, 24-29 St Giles’, Oxford, OX1 3LB United Kingdom

**Keywords:** T cell receptor, Repertoire, Paired Sequences, Alpha, Beta, Coherence, Language Model

## Abstract

T cell activation is governed through T cell receptors (TCRs), heterodimers of two sequence-variable chains (often an alpha [*α*] and beta [*β*] chain) that recognise linear antigen fragments presented on the cell surfaces. Early sequencing technologies limited the study of immune repertoire TCRs to unpaired transcripts, leading to extensive analysis of *β*-chain data alone as its greater sequence diversity suggested it should dominate antigen recognition. Over time, structural data has revealed that both *α* and *β* chains contribute to binding most antigens and highthroughput single-cell handling technologies have been increasingly applied to obtain samples of complete TCR variable region sequences from repertoires. Despite this, there is currently no repository dedicated to the curation of publicly available paired TCR sequence data. We have addressed this gap by creating the Observed T cell receptor Space (OTS) database, a source of consistently processed and annotated, full-length, paired-chain TCR sequencing data from 50 studies and at least 75 individuals. Currently, OTS contains 5.35M redundant (1.63M nonredundant) predominantly human TCR sequences and, based on recent data availability trends, will grow rapidly. We perform an initial analysis of OTS, leading to the identification of pairing biases, public TCRs, and distinct chain coherence patterns relative to antibodies. We also harness the data to build a publicly available paired-chain TCR language model, providing paired embedding representations and a method for residue in-filling that is conditional on the partner chain. OTS will be updated and maintained as a central community resource and is freely downloadable and available as a web application at *https://opig.stats.ox.ac.uk/webapps/ots*.

## Introduction

T cells are an essential component of the adaptive immune system, whose role is to selectively recognise pathogenic molecular signals from cells and effect direct killing or stimulation of the humoral (antibody) response (1). They recognise foreign entities through T cell receptors (TCRs), heteromeric pairs of sequence variable polypeptide chains. Collectively, as a repertoire of cells each with distinct TCRs, T cells can recognise millions of diverse pathogen-associated motifs.

The most studied human TCRs comprise a beta chain (βchain) and an alpha chain (α-chain), produced by genetically analogous mechanisms to the heavy (VDJ gene recombination) and light (VJ gene recombination) chains of antibodies, respectively. The most common target class for these TCRs are intracellularly or extracellularly derived, linear fragments of peptides displayed by one of six Class I or twelve Class II Major Histocompatibility Complex proteins (pMHCs) (2), which themselves are highly polymorphic across populations (3, 4). This contrasts with the contiguous molecular surfaces of extracellular proteins recognised by antibodies (5, 6). Both immunoglobulins achieve antigen binding through a combination of six Complementary-Determining Region (CDR) loops, three per chain, with the third CDR (CDR3) being the most inherently diverse as it is encoded across the gene recombination junction(s) (7).

To focus recognition on the site of variability across pMHCs, TCRs have evolved (8) and been trained during thymic development (9) to orient their hypervariable CDR3s directly above the peptide, providing maximum sensitivity to differences in antigen sequence and structure that could distinguish an errant cellular phenotype from a healthy one (10). Binding and signalling tends to be driven by energetic hotspots in the peptide, with the surrounding MHC contacted predominantly by the V gene templated CDR1 and CDR2 loops that provide sufficient complementarity to enable canonical pMHC engagement (11–13). However, despite this more general role, the CDR1 and CDR2 loops are also key in recognising certain peptides (14, 15). To avoid over-optimisation of complementarity between the TCR and the MHC, and thus a loss in specificity for the peptide, TCRs do not undergo somatic hypermutation, meaning the totality of repertoire diversity originates from gene and junction recombination diversity. Once again this contrasts with antibodies, in which any CDR can provide energetic hotspots and be tailored in sequence for the antigen during affinity maturation, generally thought to lead to increases in antigen specificity (16).

As the diverse CDR sequences across antibody/B cell and T cell repertoires hold the key to understanding individuals’ immune history, and provide opening gambits for nature-inspired drug discovery (17), much effort has been dedicated to developing V(D)J transcript amplification and sequencing approaches that can sample the diversity of receptors present in an individual at a given time (18, 19).

Early approaches involved the bulk lysis of selected cells, before preparation of sequencing libraries based on messenger RNA/genomic DNA, performing rapid sequencing (e.g. Illumina polymerase chain reactions), and processing the resulting reads (19). Such strategies provide deep samples of immune receptor repertoires (up to the order of 10^7^ reads) but are unable to capture complete B cell receptor (BCR) or TCR variable regions due to the loss of pairing information. Biases quickly emerged towards sequencing only the heavy antibody chain/β TCR chain, and particularly the CDRH3/CDRβ3 sequence, due to it being the locus of maximum diversity across receptors. There are several databases dedicated to collecting this data (20–23), and together they have spawned a broad diversity of downstream adaptive immune repertoire research and methods development (24–30).

Improvements in cell handling techniques (such as microfluidics approaches) have enabled higher throughput singlecell sequencing, now a foundational biotechnology for adaptive immune repertoire analysis (19, 31). Transcripts are tagged with a bespoke cellular barcode that enables the postsequencing reassembly of heavy/light or β/α chain pairs, albeit at a lower sequencing depth relative to bulk lysis sequencing (currently up to c. 10^5^ reads with sequencers such as 10x Genomics’ Chromium).

To study this growing body of paired-chain BCR repertoire samples, we recently built the Paired Observed Antibody Space (OAS) database and web application (32). Our curated data comprising over 2 million sequences from 12 independent studies, has enabled studies on paired-chain repertoire mining (33), structural profiling (34), and biophysical characterisation (35), as well as paired-chain antibody language modelling (36).

However, no dedicated database has yet been built to curate the rapidly emerging paired TCR sequencing data from analogous repertoire studies. The most readily accessible source of paired-chain TCR sequence data from different sources is VDJdb (37), a set of around 30,000 (sequence redundant) antigen-associated complete TCR sequences (38). While a small set of 10x-sequenced repertoires from studies between 2018-2020 have been added to iReceptor since its publication (21), they are not easily findable and require additional computational post-processing to pair up the TRA and TRB reads based on barcode identifiers.

Here, we present the Observed T-cell Receptor Space (OTS) database, which currently contains over 5.3 million paired-chain TCR reads across 50 studies. Like Paired OAS (32), all raw sequencing data is consistently processed and curated into metadata blocks to enable bespoke dataset generation, and is free to explore *via* a web application. We show how this collection of curated data reveals novel insights into TCR biology, including the identification of hitherto unidentified public TCRs and characteristic chain coherence patterns. Finally, we use OTS as the basis for building the first paired, full-chain TCR language model released to the community, based on an ESM-2 architecture (36, 39). OTS will be updated as new studies emerge, providing a central community resource for immunoinformatics studies across repertoires of complete TCR variable regions.

## Results

### Curating a database of paired-chain TCR repertoires

To collect paired TCR sequencing data, we analysed a corpus of studies that use 10X sequencing to profile immune cells, prioritising studies involving humans (see Methods). This provided an initial bibliography of 592 studies, which, when plotted by year of publication, have been steadily increasing in abundance over the past six years (Fig. 1, red dashed line). We then manually inspected these studies, looking for those which had deposited raw 10x V(D)J sequencing reads with unrestricted access on the Sequence Read Archive (SRA). Data from the resulting subset of studies were then passed through our processing pipeline, yielding 50 studies whose repertoires could be successfully processed (Fig. 1, solid blue line).

**Fig. 1.**
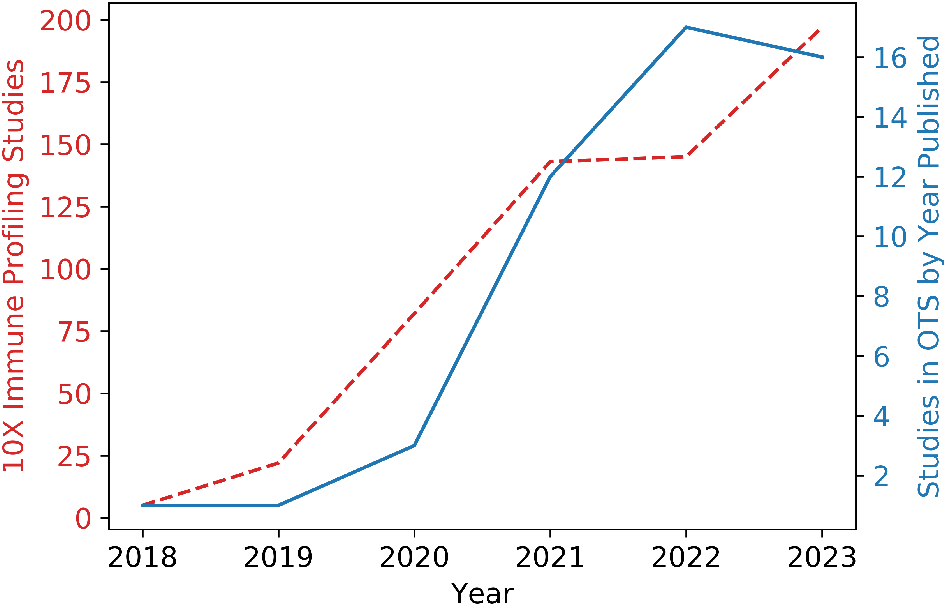
Line plots showing the growth in single cell human immune profiling data (red dashed) and public single cell human TCR repertoire data (blue solid) over time. Note that only studies published by 30/09/23 are included in the OTS figures for 2023.

Our processing pipeline was modified from that used to curate paired antibody sequencing data for the OAS database (32); full details are available in the Methods. Briefly, studies with raw fastq files deposited in the SRA were downloaded and reanalysed using 10x Genomics’ CellRanger software. Individual sequences were analysed and filtered using IgBLAST (40) and ANARCI (41) numbering, to ensure reads were full-length and free from ambiguous amino acids. Finally, pairing was carried out using the 10x barcodes.

Metadata were manually curated using both SRA-deposited information and inspection of the corresponding methods sections of each manuscript. From the 50 studies successfully processed by our pipeline, we obtained 5.3M paired sequences, which collapsed to 1.6M sequence non-redundant pairs. Redundancy arises from the sharing of sequences by expanded clones, longitudinal analysis of individuals, and commonality across individuals. Redundant sequences were retained as they may represent biologically significant clonal expansions and frequencies that relate to the metadata, with the intention that this redundancy can be collapsed for applications that require de-duplicated data.

### Diversity of datasets in OTS

The most common TCR repertoires in OTS are those from patients with COVID-19, followed by those from patients with cancer and then healthy donors (Fig. 2a). Cell types in the database are dominated by bulk (unsorted/CD3^+^/CD45^+^) T cells from peripheral blood monocytes (PBMCs), however there are also a diverse range of samples from tumour and normal tissue, as well as lymph nodes (Fig. 2b). This abundance of more general (*i.e*. not antigen-sorted) T cells might render OTS particularly suitable for investigations of the underlying grammar/language of TCRs.

**Fig. 2.**
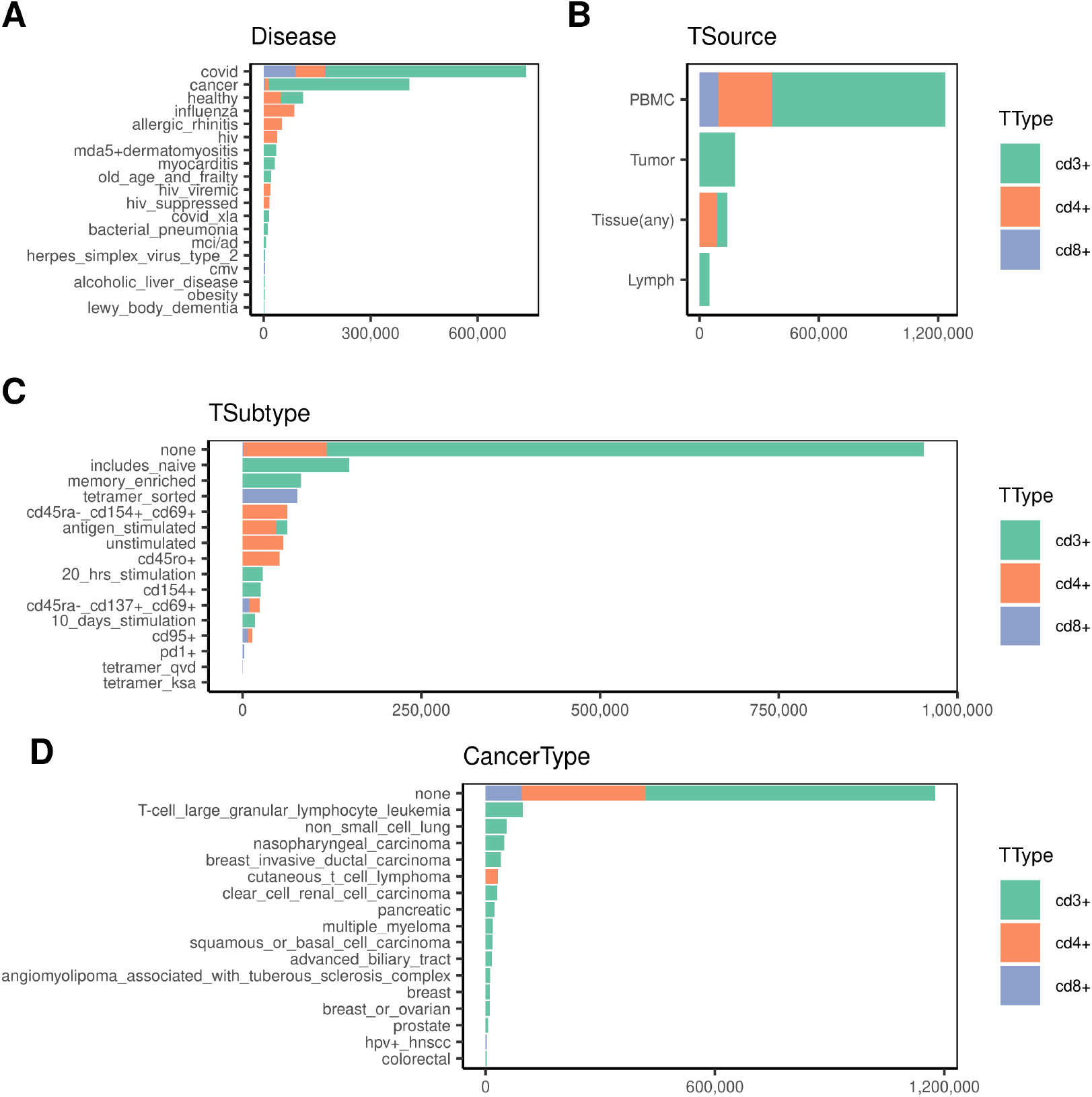
OTS covers multiple diseases, tissues, T cell types and subtypes. Non redundant paired sequence counts were divided according to the meta data associated with each sample. Counts are shown for (A) disease indication, (B) tissue source ‘TSource’, (C) T cell expansion or sorting strategy ‘TSubtype’ and (D) cancer type. Bars are coloured by CD3+/CD4+/CD8+ based on the sorting or enrichment method used to collect each sample. Where a method indicated ‘unsorted’, ‘CD45+’ or ‘live’ then these have been grouped under ‘CD3+’ for completeness. The plot titles and axis labels match the meta data and field names given on the OTS web site.

Some studies in OTS collected T cells enriched for specific memory/naïve populations, activation markers, tetramer binding or performed deliberate peptide stimulations (Fig. 2c). Collectively these constitute 40% of the total nonredundant data.

Overall, surveying the repertoire metadata in OTS allows the identification of gaps in the publicly available repertoire sample space. For example, while general datasets of CD4^+^ cells are present (mainly positively selected via magnetic enrichment methods), no study in OTS as yet solely contains gen-eral CD8^+^ T cells that were not further sorted using pMHC tetramers or surface activation markers.

### The OTS web portal and summary downloads

OTS is available as a web application from *https://opig.stats.ox.ac.uk/webapps/ots*. It can be searched by multiple parameters, such as cell surface markers and antigen specificity enrichment methods, as well as by tissue and disease state (SI Figs. 3-4). All files, including redundant sequences and full metadata can be downloaded. OTS will be periodically updated with relevant studies that both increase the depth and diversity of data available.

In the next section of the paper we explore basic immunological patterns present in OTS as a paired-chain TCR sequence resource.

### Paired gene usages indicate subtle pairing preferences related to semi-invariant TCRs and antigen specificity

We first analysed V-allele pairing in non-redundant sequences to ascertain whether any biases were evident that were not expected based on distributions of unpaired data. We estimated an expected V-allele pairing distribution by taking the outer product of the TRAV and TRBV allele frequencies in OTS in isolation. This distribution was then compared against the observed distribution in paired data (Fig. 3A-B). Over 97 % of V-allele pairings were observed at less than 0.020 % away from their expected frequency, however a number of pairings exhibited differences which were more than 6 standard deviations from the mean (Fig. 3C, Table 1).

**Fig. 3.**
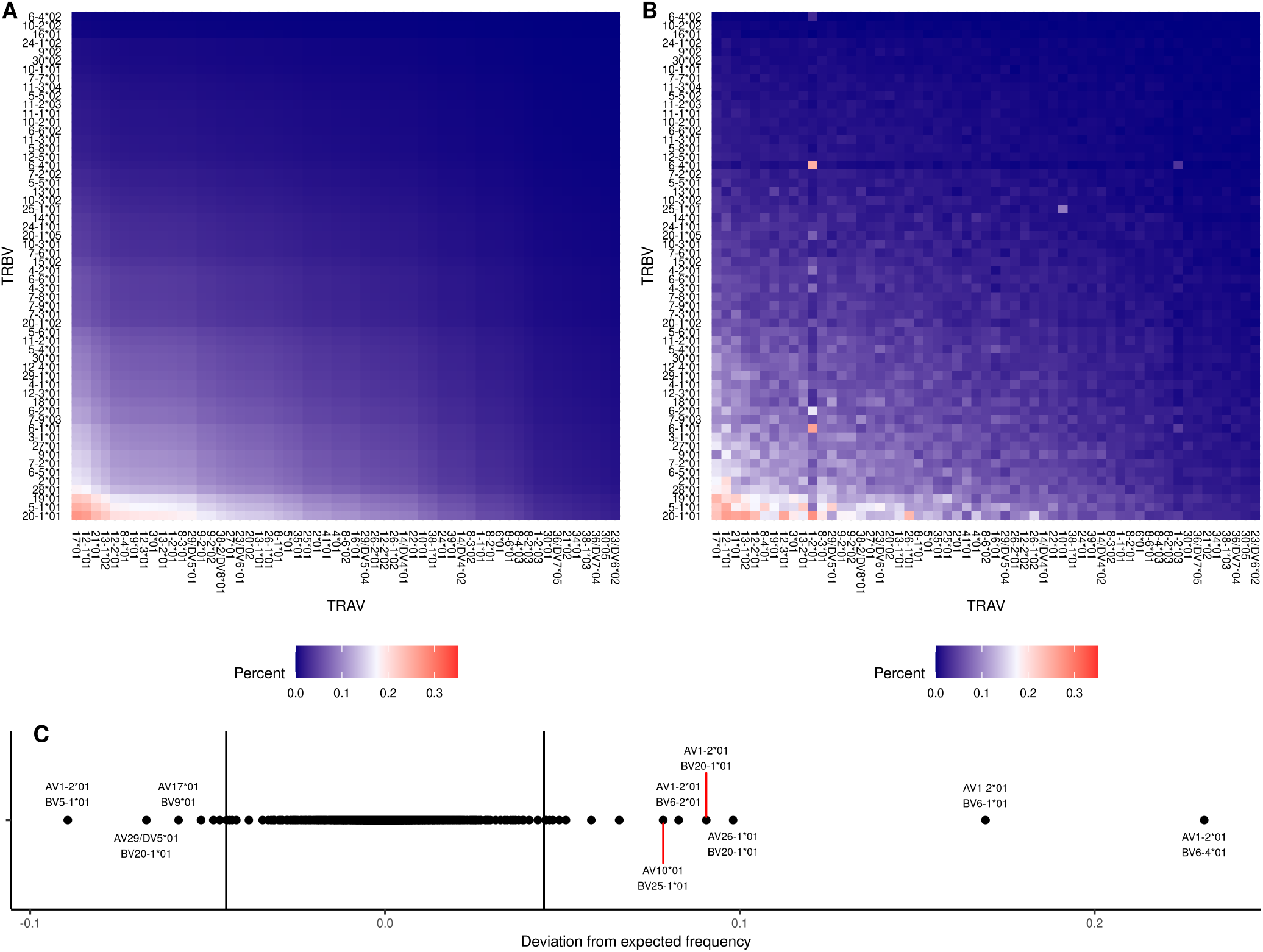
Paired sequencing data gene usage identifies unexpected pairings in bulk PBMC data linked to unconventional T cells. (A) The raw unpaired frequencies of V gene alleles for TRAV and TRBV were multiplied to form an expected counts matrix. Only the V gene alleles with top 75% highest observed unpaired frequencies are shown, frequencies were calculated for PBMC derived samples that had not been subject to further sorting (TSubtype == “none”). (B) The corresponding observed allele pairing frequencies in the same samples are shown with the same axes. (C) The expected frequencies are plotted with vertical lines indicating the 6 standard deviation cut off used to identify allele pairings that deviated highly from expected.

**Table 1.**
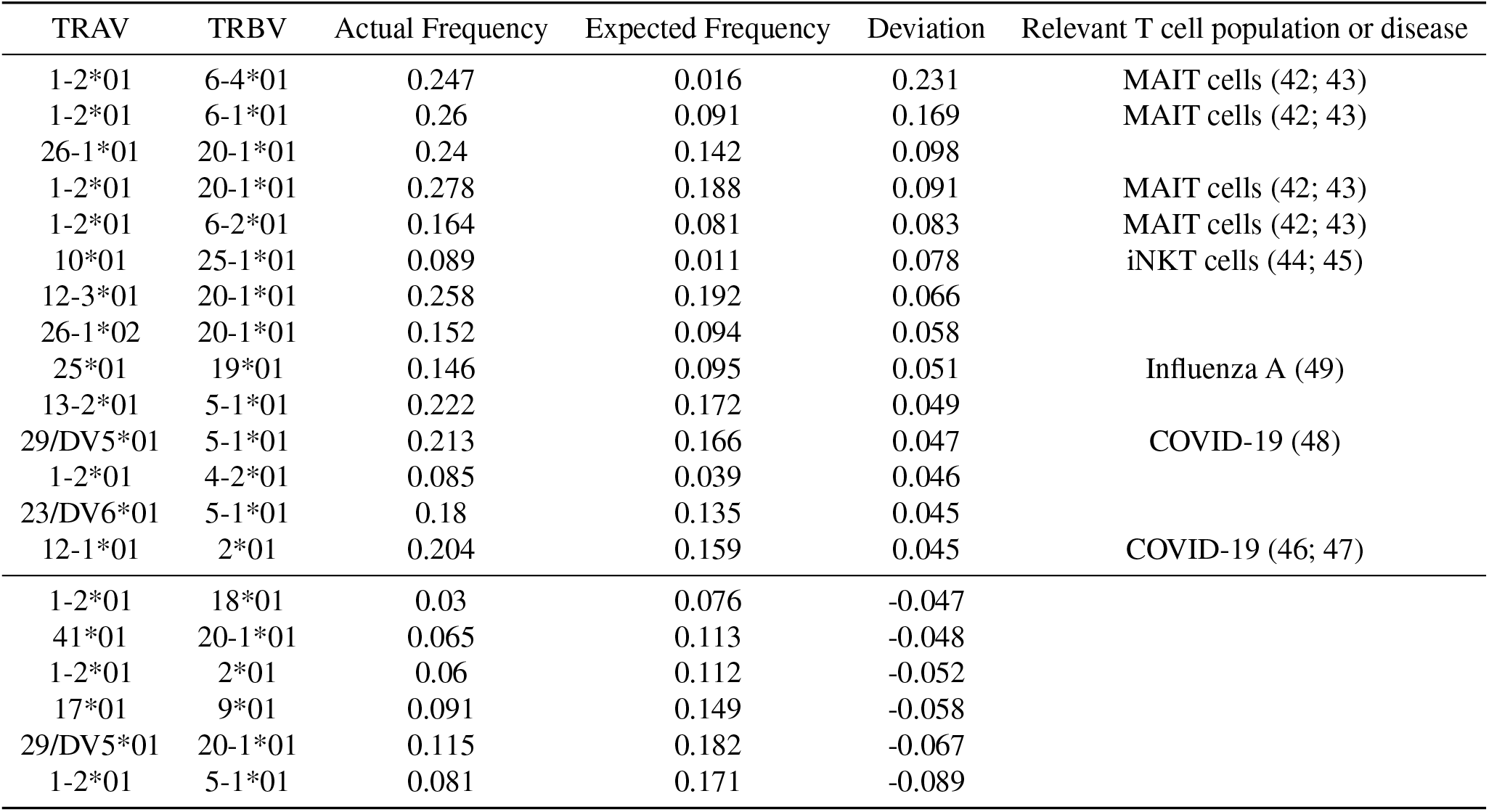
TRAV:TRBV pairings with observed frequencies over 6 SD from expected: All pairings which deviated from the expected frequency by more than 6 standard deviations are listed in the table. The corresponding literature which identify these gene usage pairings as being enriched in a specific disease indication or subset of T cells is given in the final column.

Some differences were attributable to Mucosal Associated Invariant T cells (MAITs), which adopt a semi-invariant TCR comprising TRAV1-2 paired with either the TRBV6 family or TRBV20-1 (42, 43). These pairs stood out as being amongst the most highly enriched above the expected frequency (Fig 3C, Table 1). Similarly, the highly enriched TRAV10:TRBV25-1 combination likely represents expanded populations of type I NKT cells, another semi-invariant population (44, 45).

The TRAV12-1:TRBV2 gene pairing has been detected repeatedly in SARS-CoV-2 seroconverted individuals (46) and is specifically associated with public T cell responses to the HLA-A*02-S_269–277_ epitope (YLQPRTFLL) (47).

TRAV29/DV5:TRBV5-1 has also been associated with the T cell response to COVID-19 vaccination (48). It is therefore consistent that both pairs were enriched in OTS (Table 1), given the disproportionate abundance of COVID-19 related studies and the fact that most data — irrespective of patient disease state — were collected after the start of the 2020 Pandemic.

Gene pairings associated with other common diseases were also enriched, such as TRAV25:TRBV19, implicated in the response to the Influenza A epitope HLA-A*02-M1_58-66_ (49).

Still other gene pairings were highly enriched, such as TRAV26-1:TRBV20-1 and TRAV13-2:TRBV5-1, for which there are no established functional associations. These combinations were enriched despite typically originating from genes more frequently found in the bulk repertoire.

Finally, pairings such as TRAV1-2:TRBV5-1 and TRAV29/DV5:TRBV20-1 were markedly underrepresented relative to the expected frequencies. Further work is needed to elucidate whether these pairings happen to pair less frequently or are deliberately deselected due to self-reactivity.

### Public paired TCR clonotypes highlight stereotypic responses and frequently observed lineages without known antigen associations

To further explore enriched TCRs that may be important at the population level, we next focused on TCR sequences found in multiple studies. We mined OTS to identify paired TCR clonotypes seen in at least two studies across all our datasets, i.e. ‘public’ TCRs (see Methods). A total of 2276 TCRs met this criterion, of which 45 were present in five or more independent investigations. As expected based on allele pairing frequencies, MAIT signatures were again highly enriched in the public clones; 8.5% of public clonotypes contained TRAV1-2 paired with TRBV6 genes or TRBV20-1. Gene pairings associated with particular antigen responses were also enriched, such as the TRAV27:TRBV19 pairing characteristic of HLA-A*02M1_58-66_ specificity, present in 0.7% of OTS public pairs.

We then matched these public pairs to VDJdb (37) antigenic annotations (see Methods). Of the 2276 shared clones, 74 paired-chain clonotype (TRAV+TRBV+100% CDRA3 + 100% CDRB3) matches were found to VDJdb. As class I and class II pMHCs have distinct topologies, it is unlikely that a TCR able to engage one class can also engage the other with high affinity. Therefore, as a test for consistency, we surveyed how often the 60 public TCRs with cell-surface marker annotations (CD8^+^, class-I restricted, or CD4^+^, class-II restricted) mirrored the MHC class associated with the matching VDJdb entry. We found complete agreement. However, when we expanded this analysis to match either of the individual chains in an OTS-public pair to an VDJdb annotation, the annotations were less consistent. Of the 147 public *β*-chain matched clonotypes, 6% had inconsistent cell surface markers in OTS, and of the 164 public *α*-chain matched clonotypes, 26% had inconsistent cell surface markers in OTS.

Further inspection of the paired-chain clonotype matches to VDJdb revealed that the top three most shared TCR pairs were all annotated by VDJdb as being specific to epitopes derived from the highly prevalent Epstein Barr virus (EBV). The TCR specific to HLA-A*02-BMLF1GLGTLVAML was found in 11 datasets, with an HLAB*08-EBNA3A-FLRGRAYGL specific TCR found in 9 datasets and HLA-A*11-EBNA3B-AVFDRKSDAK found in 7. The remaining shared TCRs were dominated by HLA-A*02-M1_58-66_ (Influenza-A), HLA-A*02-pp65NLVPMVATV (Cytomegalovirus, CMV) and COVID-19 specific sequences.

Most of our public TCRs (1721/2276; 75.6%) were not found in VDJdb, either as individual-chain or paired-chain clonotype matches. Of the 205 uncharacterised (‘orphan’) TCRs found in 3 or more studies, 90 were found in CD4^+^ samples, while only 36 were found in CD8^+^ samples, and 80 were found in cells with the more generic CD3^+^ label. This bias towards uncharacterised public class-II specific TCRs is likely reflective of a data bias within VDJdb towards CD8^+^ T cells and the comparative lack of studies on CD4^+^ antigenic specificity. We then further filtered our unmatched public TCRs by those whose observed V gene pairing frequencies were lower than the median value (0.0226 %) and whose CDR3 lengths were greater than the respective median values (CDRA3: 12, CDRB3: 12). This highlighted nine clones unlikely to be public by chance (Table 2). These results show how OTS can be used to identify understudied or unexpected public TCRs that may be carrying out important protective functions.

**Table 2.**
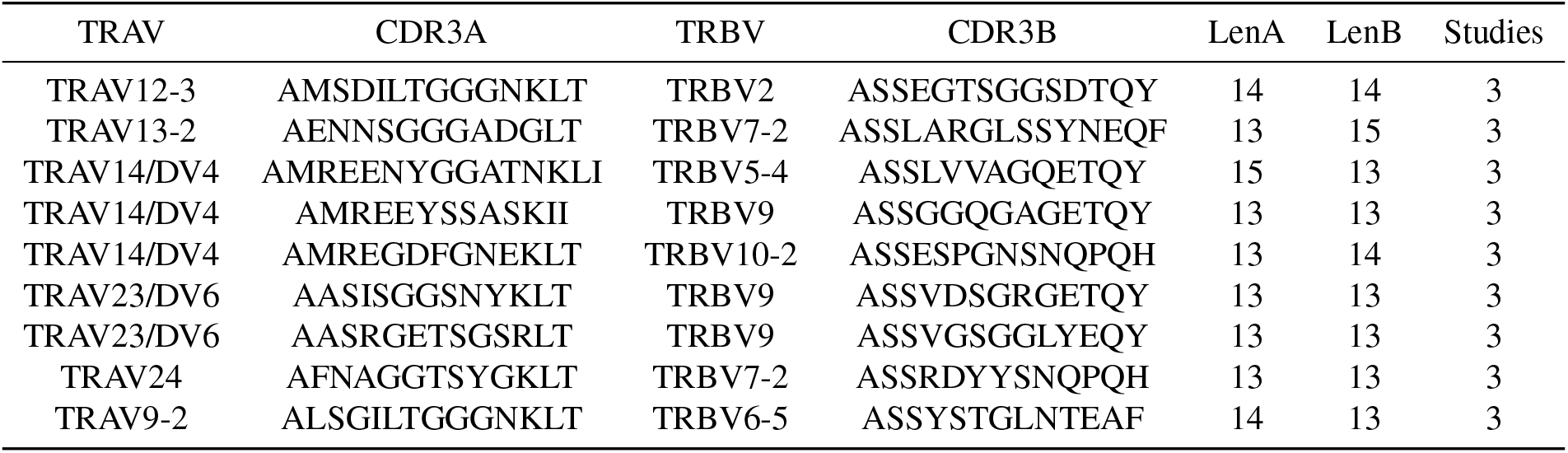
Public TCRs found in OTS which had longer than median CDR3 sequences, did not arise from highly frequent V gene pairings and were not found in VDJdb: TCR sequence pairs defined as publinc in OTS (i.e. present in multiple distinct studies) with no single chain overlap in VDJdb (V gene and CDR3 sequence in either chain), were filtered by CDR3 length greater than the median value for each chain (12), and for those V gene pairings which were present at less than the median observed pairing frequency in bulk PBMC data (values taken from the observed statistics detailed in Fig. 3 and previous sections).

### Antigen-selected TCRs exhibit more balanced chain coherence patterns than memory BCRs

A recent study by Jaffe *et al*. revealed that memory BCRs exhibit a phenomenon they termed ‘light chain coherence’ across individuals (51). That is, when a memory BCR heavy chain clonotype is found in different individuals (is public), it is commonly found (*∼* 80% of the time) accompanied by the same light chain V gene. The phenomenon does not apply in reverse; public light chain clonotypes found in the memory populations of different individuals are uncommonly (*∼* 6% of the time) found paired with the same heavy chain V gene. Neither chain displayed a coherence signal in the naïve BCR populations, suggesting that minimal intrinsic pairing biases exist and that the phenomenon arises only in the context of ‘functional’ BCRs selected against common antigens. The implication is that the light chain V gene identity may play a key role in correctly orienting the heavy chain CDRs for antigen binding (52), or contributing key germline interactions *via* its own CDR loops (5, 6, 53).

As the energetics of TCR:pMHC complexes appear more evenly shared between the *β* and *α*-chain than do the energetics of antibody:antigen complexes between the heavy and light chain (13), it is likely that any coherence signal would also be more balanced between chains for TCRs. We used OTS to perform an analogous, contrastive investigation to Jaffe *et al*. on TCR repertoires.

First, we curated a set of TCR repertoires to act as a baseline for chain coherence values (‘baseline’), and a separate set enriched for common functions on the basis of tetramer sorting (‘antigen sorted’) (54–56). This baseline set is not a purely naïve TCR dataset, and contains a mixture of cell types; this approach was necessary due to the paucity of studies that sort for naïve peripheral T cells. We then calculated *α*-chain coherence on *β*-clonotypes and *β*-chain coherence on *α*-clonotypes, based on an initial clonotype definition of a matching V gene and 100% CDR3 sequence identity and matching partner V gene (Table 3, see Methods).

**Table 3.** Mean *α*- and *β*-chain coherence values for the Baseline and Tetramer datasets derived from OTS. Redundant statistics are based on a dataframe that allows identical paired sequences to be found in different repertoires from the same study; these are filtered out to retrieve the non-redundant statistics. CDR3 Levenshtein Distance (LD) reflects the maximum number of edit distances between CDR3 sequences belonging to a clonotype (50). Clonotypes and partner chain coherence are defined at the level of the gene.

| Redundant/Non-Redundant | CDR3 LD | Category | Mean α-chain Coherence | Mean β-chain Coherence |
| --- | --- | --- | --- | --- |
| Redundant | 0 | Baseline | **9.55%** (N=33868) | **4.92%** (N=152998) |
|  | 1 | Baseline | 4.78% (N=19415) | 3.68% (N=41654) |
|  | 2 | Baseline | 3.44% (N=9429) | 3.31% (N=12583) |
|  | 0 | Tetramer | **45.59%** (N=124) | **41.54%** (N=874) |
|  | 1 | Tetramer | 43.21% (N=110) | 25.52% (N=700) |
|  | 2 | Tetramer | 41.78% (N=104) | 20.43% (N=574) |
| Non-redundant | 0 | Baseline | **9.15%** (N=33829) | **4.90%** (N=152969) |
|  | 1 | Baseline | 4.68% (N=19377) | 3.67% (N=41638) |
|  | 2 | Baseline | 3.42% (N=9390) | 3.31% (N=12557) |
|  | 0 | Tetramer | **17.74%** (N=95) | **11.55%** (N=811) |
|  | 1 | Tetramer | 16.16% (N=85) | 9.16% (N=653) |
|  | 2 | Tetramer | 13.26% (N=81) | 7.21% (N=540) |

Baseline mean coherence values for both the *α*and *β*-chains were low and roughly equivalent (9.55% and 4.92%; *N*=33868 public *β*-clonotypes, *N*=152998 public *α*clonotypes, respectively), consistent with the values seen for naïve antibodies. However, contrary to observations in antibodies, both public antigen-sorted *α*and *β*-chains exhibited significantly more coherence, and by similar amounts (45.59% and 41.54%; *N*=124 public *β*-clonotypes, *N*=874 public *α*-clonotypes respectively). This is consistent with either or both chains being able to dictate pMHC specificity. The lower absolute coherence for the *β*-chain could suggest that the same *β*-clonotype might more frequently associate with *≥* 2 antigens than in antibodies, or reflect the fact that the germline-encoded CDRs in TCRs are more promiscuous, leading to antigens on average imposing a weaker selection on the V genes.

We assessed the impact of making the coherence criteria stricter, by defining clonotypes and matching genes at the level of the allele (SI Table 1). This typically led to modest decreases in coherence values but analogous trends were observed between Baseline and Tetramer-sorted populations. We also measured the impact of relaxing the clonotype definition to allow CDR3s within a Levenshtein distance (LD) of 1 or 2 and a matching V gene to be grouped together (Table 3). This again led to drops in coherence statistics across all categories, but yielded similar patterns.

One potential confounding factor in calculating the extent of coherence is that the same sequence could be recorded as being found in multiple individuals from a single study, but in fact was observed frequently only due to contamination in the sequencer. Therefore, we sought to obtain a lower bound for the coherence signals by removing instances where the same *αβ* sequence is observed amongst multiple individuals in the same study. This led to baseline *α*and *β*-chain coherence values of 9.15% (*N*=33829 public *β*-clonotypes) and 4.90% (*N*=152969 public *α*-clonotypes), and antigen-sorted *α*and *β*-chain coherence values of 17.74% (*N*=95 public *β*-clonotypes) and 11.55% (*N*=811 public *α*-clonotypes), respectively. A larger reduction in coherence signals was observed amongst antigen-selected TCRs on applying redundancy filtering than was observed for the light chains of memory BCRs (51), but this is unsurprising due to the lack of somatic hypermutation in TCRs, meaning it is likely that more genuine signal is being filtered out.

### Paired-chain TCR language model

Finally we used the data contained in OTS to build a publicly available paired TCR language model (“TCRLang-Paired”). Using the existing AbLang2 architecture (36), we started by training on over 9M curated non-redundant, unpaired sequences from iReceptor (4.58M *α*-chains, 4.62M *β*-chains) using the masked residue prediction task (see Methods). The model was then finetuned for a smaller number of steps on around 1.35M non-redundant OTS pairs. We evaluated the model’s performance after training with two different loss functions: cross entropy loss (CEL), where all mistakes are penalised equally, and focal loss (FL), which rewards a model more for predicting rare phenomena accurately (36).

First, to explore the embedding space of the two models, we generated sequence embeddings of all non redundant paired TCRs sequences present in OTS and visualised them using tSNE plots (Fig. 4a-b, SI Fig. 1A-F). Each data point represented a paired *αβ* TCR sequence and could therefore be coloured by one of 4 genes (TRAV, TRAJ, TRBV or TRBJ). For the model trained with FL, annotation by TRAV gene most cleanly defined all dominant clusters on the plot (Fig. 4a). In contrast, the clusters derived from the CEL model were best defined by TRAJ gene annotation (Fig. 4b). For each model the remaining genes were also visible as sub clusters dispersed within the larger areas of density (SI Fig. 1A-F). These results suggested that both models were learning sensible paired representations of TCRs which related to their variable regions. They also highlighted the fact that varying the loss function can teach a model to focus on different biological features within the same class-specific context. In general sequence recovery tasks, the model trained with CEL performed marginally better, so this was chosen for all downstream tests and as the final basis for TCRLang-Paired.

**Fig. 4.**
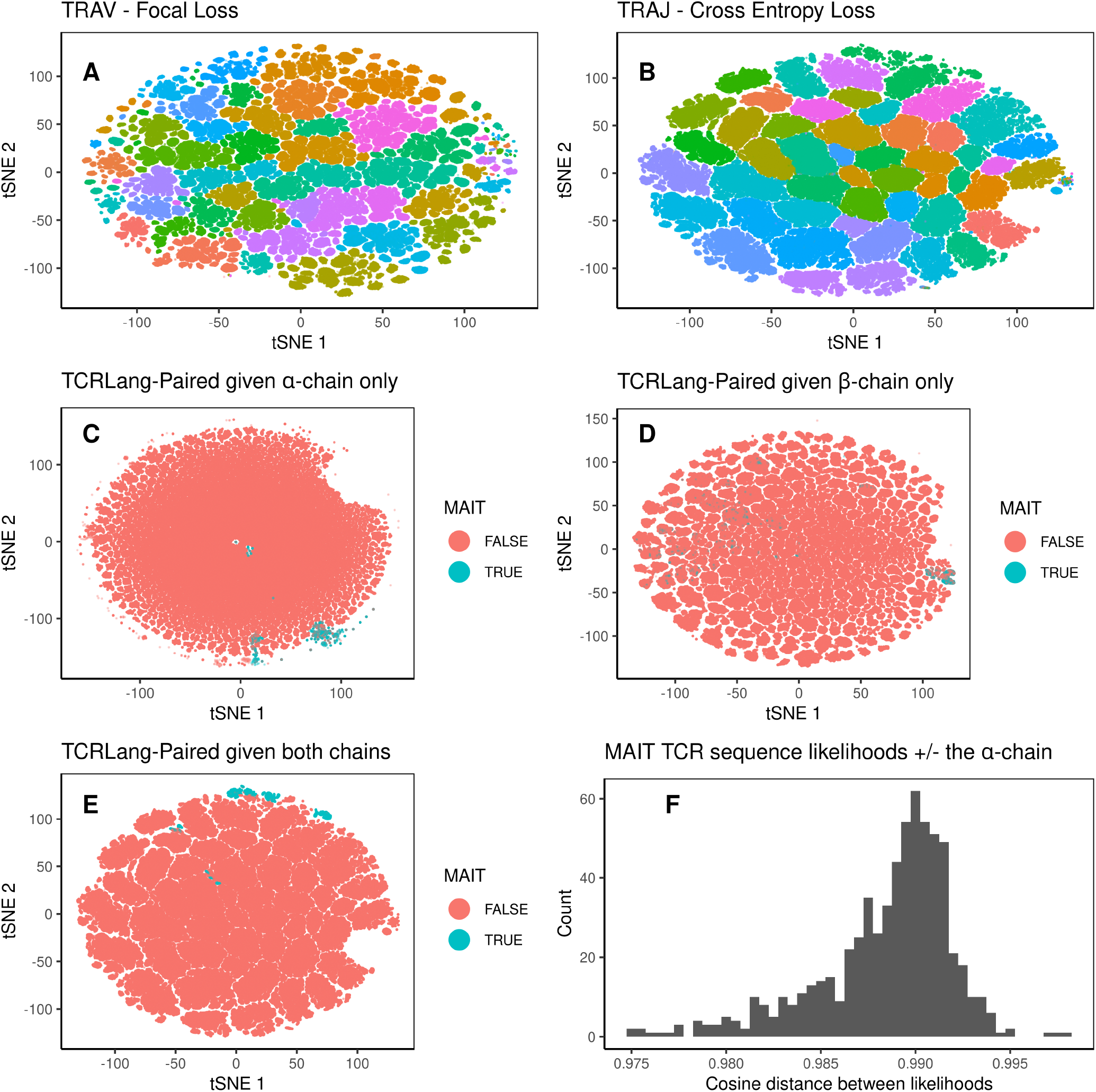
TCRLang-paired can distinguish TCRs based on V/J gene usage and group unconventional subsets in similar regions of embedding space. TCRLangpaired embeddings were generated for all non redundant *αβ* TCR pairs in OTS. The paired embedding space was visualised with tSNE analysis for a model trained with focal loss (FL) and a model trained with cross entropy loss (CEL). The two best separated representations for each model are shown, with points coloured by (A) TRAV gene usage for FL and (B) TRAJ gene usage for CEL. Remaining plots show the embedding space of the final model, TCRLang-paired, trained using CEL. For this model, (C) the *α*-only, (D) *β*-only, and (E) paired embedding spaces were also coloured by whether TCRs had the MAIT cell signature or not. (F) The cosine similarity distances between residue likelihoods of the MAIT TCR sequences generated using information from both chains or the *β*-chain alone are shown.

To further test whether the embedding representation takes into account information from both chains, we looked at sequences associated with MAIT cells (characterised by specific TRAV, TRAJ and TRBV gene usage). We categorised all data points based on whether or not they expressed the MAIT-associated gene usage combination (see Methods). We then used this information to annotate data points on tSNE plots created from embeddings generated from either the *α*-chains alone (Fig. 4c), the *β*-chains alone (Fig. 4d), or the paired sequences (Fig. 4e). The resulting plots showed that this functionally constrained population of TCRs were more co-localised in tSNE space for the paired representation, while for the unpaired representations, particularly for *β*-only, MAITs were spread more diffusely with less of a separation from the background distribution of non-MAIT cells.

We next tested the ability of the model to infill missing residues (sequence recovery) in the CDR3 of the *α*-chain, when given only that chain (equivalent functionality to the first AbLang model (28)), or when the partner *β*-chain was also provided (equivalent functionality to AbLang2 (36)). We continued to focus on MAIT TCRs and processed all sequences from a separate study by Garner *et al*. which focused on MAIT cells (57) and selected only those sequences which were not already present in OTS. However, when asked to infill the entire missing CDRA3 region of these unseen sequences, the context of the *β*-chain did not improve sequence recovery and gave slightly worse results (SI Fig. 2, Levenshtein distance (LD) ≤ 1: paired recovery 46.8% vs unpaired 48.0%). When we flipped the task to predict an entire masked CDRB3 sequence, the recovery values were worse (SI Fig. 2, LD *≤* 1: paired recovery 1.1% vs unpaired 1.8%), however the unpaired predictions were still marginally better.

Given the difficulty of the masking task, we decided to perform a simpler experiment to assess whether TCRLang-Paired was incorporating information from the *α*-chain into the representation of the *β*-chain (or visa-versa). Again using unseen TCRs from Garner *et al*., we generated likelihood matrices for each TCR of the *β*-chain with, and without, the sequence of the partner *α*-chain. We then calculated the cosine similarity distance value for each pair of matrices. While similar, no pair of likelihood matrices were exactly the same, with the distribution of cosine similarity distances ranging from 0.975 to 0.999 (Fig. 4f). This confirmed that the MAIT *β*-chain representations differed depending on the presence of *α*-chain information. Despite limitations in predicting masked residues, these analyses do suggest that both intra-chain and inter-chain features are learned by TCRLang-paired, which might reasonably assist downstream prediction tasks that depend on complete binding site features.

## Discussion

The Observed Antibody Space (OAS) databases (20, 32) are now established resources in the field, enabling the generation of models for tasks such as therapeutic humanisation (24, 27), repertoire mining (30, 33), repertoire structural profiling (34, 35) and large language modelling (26, 28, 29, 36). Here, we provide the first database for analogous explo-rations of paired-chain TCR repertoires, which we hope will stimulate/support analogous research efforts.

We collected data from 50 studies investigating viral infections, cancer and autoimmunity, including a broad range of tissues and T cell subtypes. This yielded over 1.6M non redundant paired TCR sequences that we explored from various perspectives.

The majority of the data in OTS came from bulk unsorted (CD3^+^) or CD4^+^ populations; CD8^+^ T cells were represented but were always further isolated into a sub-type that may have reflected memory/activation status or antigen specificity. The large quantity of bulk unsorted data allowed us to perform analyses on V gene pairing biases, particularly in the deviation from expected frequencies calculated from the individual chains. While the majority of TRAV:TRBV pairings fell close to their expected frequencies, we found several which deviated. Some of these could be accounted for by pairing preferences of semi-invariant TCRs specific to MR1 (MAIT cells) and CD1d (iNK T cells), while others were likely associated with public responses to COVID-19 epitopes in highly prevalent HLA alleles such as HLA-A*02. However, some pairing deviations, both above and below expected frequencies, could not be accounted for and may warrant further investigation. As the volume of OTS data rises, more and clearer patterns could emerge.

We searched for the presence of public TCRs across studies in OTS, using detection in independent studies to ensure overlap was not a result of contamination. We found 2276 public TCR sequences and were able to annotate the pMHC reactivity of several of them through a comparison to the clones in VDJdb (37). The most highly shared sequences were specific to immunodominant epitopes from globally prevalent viruses such as EBV, Cytomegalovirus, COVID-19 and Influenza-A, presented on common alleles such as HLAA*02 and HLA-B*08. T cell immunology has a historic focus on CD8^+^ T cells and has skewed towards alleles which are more prevalent in Caucasian populations. Therefore, public sequences and corresponding specificity annotations are more likely to exist for TCRs which see immunodominant viral epitopes presented on certain HLA alleles. However we found many more public TCRs that had no annotation in VDJdb, but that were nonetheless highly shared and were not the result of highly probable recombination events (based on their usage of less common V gene pairings and longer CDR3 loops). These sequences are likely to have explicable functional origins, which may become understood in time as high-throughput pMHC assay technology develops and as repertoires from a broader diversity of HLA alleles are studied.

In our comparisons of overlap with VDJdb, we observed that our T cell population annotations (CD4^+^ or CD8^+^) showed 100% agreement with the given specificity (MHC class I or class II) for paired chains. This is consistent with the topology across the TCR variable region being complementary to the peptide presented by one or the other class, but not to both. We expanded this to consider single-chain overlap with VDJdb, and found that the level of agreement dropped.

One possible explanation for this is that methods such MACS sorting, which are used to enrich for bulk populations, have a non-negligible error rate, and that, while the bulk sorted population is predominantly CD4^+^, some CD8^+^ T cells will have filtered through (e.g. double positive cells). Both of these effects reduce the reliability of interpreting the surface marker annotation as reflective of the MHC restriction of all the TCRs in the repertoire sample. However it is also likely that, at some level, this loss in accuracy corresponds with the importance of considering both chains to accurately infer the specificity of a TCR. That is, MHC class specificity, let alone peptide specificity, can be less reliably assigned based on a single-chain clonotype match, even at the level of matching V gene/100% CDR3 identity.

Our study on the single-chain clonotype:partner V gene coherence of 10x-sequenced, antigen-sorted TCRs demonstrates that an enhanced coherence signal over baseline is detectable for both chains. The degree of enrichment over baseline pairing preferences remains an estimate, with values varying depending on the stringency of sequence nonredundancy filtering and clonotype definition, and with precision limited by the lack of a true naïve TCR baseline and the presence of only three antigen-sorted studies sorted TCR sequences (54–56), not all of which were tetramer-sorted for the same antigens. The number of public clonotypes across the antigen-sorted data is also relatively small. As such, we anticipate that these coherence values will converge as new studies are added to OTS. The relative coherence of *α*- and *β*chains is considerably more even than the balance observed for memory BCRs, where the light V gene exhibits extreme coherence on the heavy chain clonotype while the heavy V gene shows no coherence on the light chain clonotype. This potentially reflects the fact that CDRB3 sequences dominate less over CDRA3 sequences in determining specificity than do CDRH3 sequences over CDRL3. This is supported by our recent work on TCR structure prediction, which highlights that CDRA3 and CDRB3 loops are as structurally diverse as one another and currently as challenging as one another to model (58). The lower value of both *α*- and *β*-chain coherence than light chain coherence may reflect the fact that the TCR V gene-encoded CDR loops play a less important role in peptide recognition than the CDRL1 and CDRL2 loops do in antigen recognition for antibodies. Higher coherence signals may be observed between different regions of the TCR sequence, as suggested by the observation of ‘simultaneous near-coincidences’ of paired CDR3 sequences in other tetramer-sorted repertoires (59).

Finally, we utilised the data in OTS to build a publicly available standalone paired-chain TCR language model (‘TCRLang-Paired’); others have studied the ‘language’ of combined paired-chain TCR:pMHCs (60, 61). TCRLang-Paired is capable of infilling missing residues from both chains as well as producing sequence embeddings which may be used for downstream analyses such as sequence clustering and zero shot predictions. The results of sequence embeddings clustering and differences in chain likelihood distributions (with and without the partner chain) demonstrated that TCRLang-Paired learns and considers information from both chains in making its predictions. In building TCRLang-Paired, we used the architecture established for AbLang2, and explored two standard large language model loss functions, as a simple proof-of-principle of the utility of OTS data. Even with this preliminary exploration, we found evidence that the choice of loss function influences which biological features are most prominent in the paired representations of a single class of immune receptor. This highlights how the training process of future language models might be rationally designed according to the intended application of the embeddings in downstream tasks.

We have not included data deriving from other paired sequencing methodologies such as CLINT-Seq, PARSE, or custom pipelines such as those used in Tanno *et al*. 2019 (62) and Spindler *et al*. 2020 (63), in OTS. As these (or new) methods begin to underpin a larger proportion of single cell studies, we will consider how data from such techniques can be incorporated into our pipeline while maintaining consistency and comparability. Additionally, although we were able to access 10x data from a large number of studies, the necessity for raw fastq files further limited which datasets could be added to OTS; many studies only supply processed data. While this is valuable, we chose not to incorporate it into our pipeline as we are unable to confidently retrieve full length sequences, neither can we be confident that the various versions of software such as CellRanger do not result in differences in the data. Other studies release the raw files but behind access limitations, which also directly prohibits their use in OTS.

Nonetheless, we have demonstrated that the repertoire samples we have already included in OTS are valuable to diverse fields, principally those of repertoire-based drug discovery, basic adaptive immunology, and immunoinformatics methods development. We hope OTS will play a role in directing further improvements in coverage by explicitly highlighting regions of public paired sequencing data paucity.

## Methods

### Literature search for Public 10X V(D)J Sequencing Data on TCRs

To obtain a body of papers enriched for studies that release paired TCR sequences, we used the publications page of the 10x Genomics website (https://www.10xgenomics.com/resources/publications), which scrapes the academic literature for studies that employ 10x sequencing. As of 30^th^ September 2023, this database contained references to over 6,000 papers. We first filtered this literature corpus for studies involving humans that performed “Single Cell Immune Profiling”. This yielded a subset of 572 studies, which we then further filtered for those that mention TCRs in the manuscript text, performed explicit paired V(D)J sequencing, and made their raw (unprocessed) data publicly available in a repository such as the Sequence Read Archive (SRA). Datasets from the resulting 50 studies were carried forward as input to the OTS read processing pipeline.

### OTS Data Processing Pipeline

The OTS processing pipeline is based on the Paired Observed Antibody Space (OAS) pipeline (32), modified to handle the distinct properties of TCRs. SRA numbers from each study were used to retrieve fastq files via the SRA toolkit version 3.0.7.

CellRanger version 7.1 was used to process and filter the raw fastq data into fasta files which contained barcoded nucleotide sequences. Parameters were adjusted to search for TCR sequences only.

The nucleotide sequences were translated and aligned using IgBLASTn version 1.17.1 (40) using human and mouse germline sequences downloaded from IMGT (64). Unaligned and nonproductive (flag: productive = False) sequences were removed. Each sequence was then IMGT numbered using ANARCI (41). Sequences that could not be processed by ANARCI were removed.

TCR-α and TCR-β chains were paired using 10x barcode information. Those barcodes which mapped to multiple α or β-chains and therefore had ambiguous pairing were removed. Meta data was compiled using both information in the SRA tables associated with each file, as well as the methods detailed in the manuscript associated with each study. This data was unified into a consistent format designed to allow users to group together relevant samples or search for specific features (for example, disease, tissue or phenotype) that may be of interest.

For two studies explicitly focused on *γδ* T cells we modified the pipeline to include these sequences. The standard CellRanger pipeline removes *γδ* chains during a filtering step. To work around this, we processed the unfiltered sequences output by CellRanger and selected only *γδ* pairs. The collected sequences were then processed and filtered in the same manner detailed above.

### Paired V-allele usage analysis

Non redundant full-length sequences by study (i.e. those which were not present more than once in all datasets from a given study) were analysed. Only those studies which did not further sort populations by cell surface marker or phenotype were included. A distribution of expected V-allele pairing frequencies was calculated using the V-allele frequencies in isolation (values were percentages of the total TCRs analysed) and taking the outer product to build a matrix. This gave an expected distribution of every possible TRAV and TRBV allele pair present in the data. These were visualised in a heatmap format (Fig. 3A). To reduce the number of data points on the heatmap, only V-alleles in the top 75% of the unpaired frequency distribution were shown, these were ordered by frequency (highest to lowest from the origin). The corresponding observed frequency distribution was extracted from the paired data (the percent of times a given V-allele pairing was observed over the total number of pairs). To calculate the deviation between observed and expected for each pairing, the expected value was subtracted from the observed value to give a positive deviation if higher than expected, and a negative deviation if lower than expected.

### Overlap with annotated TCRs from VDJdb

All human sequences (56664 paired and 84388 unpaired) were downloaded from VDJdb (https://vdjdb.cdr3.net/, December-2023, (37)). These were filtered to remove commercial data (those depositions not associated with an academic study) leaving 14642 paired and 42173 unpaired sequences. Paired overlap with data in OTS was analysed by matching both TRAV- and TRBV genes and exact CDRA3 and CDRB3 sequences for a pair. Unpaired overlap just considered a match in either chain of the V gene and CDR3.

### Chain Coherence Analysis

The degree to which partner chain V genes cohere within public single-chain TCR clonotypes was explored using a similar protocol to Jaffe *et al*. in their study on BCRs (51). To illustrate the pipeline, average **alpha** chain coherence was evaluated by the following protocol:

1) Cluster the data into **beta**-clonotypes (β-clonotypes), defined as TCRs deriving from the same TRBV allele and having within a threshold CDRB3 amino acid sequence identity.

2) Filter to preserve only β-clonotypes seen in at least two individuals. Studies for which we could not map repertoire samples to individuals were treated as one large pseudo-repertoire deriving from a single individual for the purposes of this analysis. Multiple instances of the same β-clonotype seen with the same TRAV allele in an individual were collapsed to a single instance, to mitigate against large clonal expansions in a single individual swamping the patterns seen across individuals.

3) Evaluate the *α*-chain coherence for each *β*-clonotype, as:

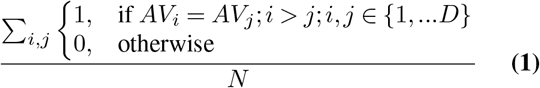

where D is the total number of alpha V (AV) gene instances, and N the total number of pairwise AV gene comparisons made within the β-clonotype, which goes with 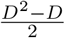. Therefore, a public β-clonotype where all partner AV genes are the same as one another would have a coherence score of 1 (100%), while one where all partner AV genes are different would score 0 (0%). A public β-clonotype associated with three AV genes of which two are the same would score 1/3 (33.3%).

4) Calculate the average *α*-chain coherence by taking the mean value across all public *β*-clonotypes.

*β*-chain coherence was evaluated in an analogous fashion, clustering the data first into public *α*-clonotypes and then calculating pairwise beta V (BV) gene identities.

In the first experiment, all 5.35 million sequences within OTS (30^th^ September, 2023) were collected into a single data frame alongside study metadata. The studies were then split into a set of tetramer-sorted TCR sequences (54–56) (‘antigen-sorted’, analogous to Jaffe *et al*’s. ‘functional’ antibodies) and a set of unsorted TCRs to offer baseline coherence values (‘baseline’, an imperfect approximation to Jaffe *et al*’s naïve B cells). Average coherence values for each TCR chain were calculated and contrasted across the two sets.

To account for the possibility of contamination across samples within a study, we also calculated average coherences based on a reduced data frame, where the sequences from each study (regardless of individual from which they were sourced) were clustered to be non-redundant at a 100% sequence identity threshold.

Clonotypes were defined as clones having a matching V gene allele and a CDR3 sequence within a Levenshtein distance (LD) (50) of 0, 1, or 2 from other sequences in OTS. For distances *≥* 1, greedy clustering was used for computational efficiency.

### Paired TCR Language Model

We first curated separate datasets of unpaired and paired TCR sequences with which to train the model.

Paired TCR sequences from OTS were first filtered to remove those deriving from tetramer-sorted datasets (54–56) or mice. The remaining TCRs were clustered at a 100% sequence identity level across the concatenated *αβ* sequence, to remove redundancy. From this set, 200K sequences were randomly sampled, of which 100K were assigned to the validation set and 100K were assigned to the test set. The remaining TCRs were shuffled and represented our training dataset of 1.36M paired sequences.

Unpaired TCR sequences were obtained from the iReceptor database (21), which, as of 30^th^ September 2023, contained around 3.64Bn *β*-chain and 880Mn *α*-chain sequencing transcripts. Full-length amino acids reads were derived by filtering iReceptor for (TRA or TRB) PCR targets and using the keyword term “complete+untemplated”. This yielded around 1.1M redundant single-chain sequences, which clustered into 404K non-redundant sequences, of which around 194K were *α*-chains and 230K were *β*-chains.

To boost the number of unpaired reads, we explored using a ‘germline in-filling’ protocol, in which we substituted truncated V gene reads for the amino acid sequence of the allele to which they were assigned (‘v_call’ column); we curated a reference set of TRAV and TRBV allele pre-CDR3 junction region amino acid sequences using the IGMT Gene-DB (64). The accuracy of this protocol (*c*. 93%) was benchmarked by comparing sequences generated by in-filling to ground truth full-length reads from the two Illumina NovaSeq 6000 datasets in iReceptor (65, 66). We then searched for (TRA or TRB) PCR targets isolated using Illumina MiSeq sequencing (sequencing platform keyword: “Illumina MiSeq”); on manual inspection, these reads were almost entirely complete in the J gene and CDR3 junction regions but were always very prematurely truncated in the V gene encoded region. This time we added the keyword filter “Healthy” to capture repertoires with a lower probability of skewing towards certain clones. All sequences were passed through our germline in-filling pipeline, leading to around 11.29M non-redundant sequences, of which 5.88M were *α*-chains and 5.40M were *β*-chains.

The two *β*-chain and *α*-chain datasets were combined according to chain type and then processed through ANARCI (41), filtering out those that could not be recognised as TCRs, that lacked the conserved disulfide bridge, that contained stop codons or ambiguous amino acid calls (e.g. residues labelled “X”), or that had a sequence length of < 102 residues (indicative of substantial truncation). Any residues outside of the IMGT-defined variable domain were deleted. Finally, we filtered for redundancy at a 100% sequence identity threshold (both internally and to either the *β*-chains or *α*-chains of the TCRs in the paired sequence test set, as appropriate). The resulting single-chain training TCRs were shuffled and comprised around 4.58M *α*-chains and 4.62M *β*-chains.

The paired TCR language model was implemented in PyTorch 2.0.1 (67) and was based on the paired antibody language model (AbLang-2) from Olsen *et al*. (36). Briefly, we trained a 12-layered version of the architecture from ESM2 model using an embedding size of 480, the SwiGLU activation function (68) and the Adam optimiser. The model was first trained on the unpaired TCR beta and *α*-chain sequences for 200,000 steps using a linear warm-up for 1,000 steps (peak learning rate: 0.0004) and a cosine learning rate decay over 199,000 steps (weight decay: 0.01). It was then fine-tuned for 10,000 steps on paired TCR beta and *α*-chain sequences (peak learning rate: 0.0001). The effective batch size was 8192 during training, with a layer normalisation rate of 0.1 and ε = 1e-12. For each batch, training was performed using cross entropy or focal loss on the task of either (a) randomly masking and predicting individual residues (selected dynamically), (b) randomly masking and predicting sets of 3-5 consecutive residues, and (c) randomly masking and predicting one large string of consecutive residues (36, 69). The proportion of masked residues per batch set was fixed at 15%.

### Model Evaluation Methods

TCRLang-Paired was used to generate embeddings of all human non-redundant paired sequences present in OTS. Sequence embeddings were generated (see: https://github.com/oxpig/AbLang2) for both chains, as well as for each chain in isolation. The embeddings were used to generate 2D tSNE coordinates used to plot the embedding space. Each datapoint represented a single paired TCR which could be coloured by gene usage or whether it belonged to the MAIT family. MAIT TCRs were identified by having V gene usage of TRAV1-2 combined with J gene usage of TRAJ20/TRAJ12/TRAJ33 in the *α*-chain, with V gene usage of TRBV20-1/TRBV6-1/TRBV6-2/TRBV6-3/TRBV6-4/TRBV6-5/TRBV6-6/TRBV6-8/TRBV6-9 in the *β*-chain. Sequence likelihoods (see: https://github.com/oxpig/AbLang2) were generated for MAIT TCRs present in Garner et al. 2023 (57). The resulting likelihoods had dimensions of [Sequence length, Tokens] and were softmaxed (for each position in the sequence), before the entire matrix was flattened to allow calculation of cosine similarity differences.

### Data Analysis and Visualisation

Data was analysed using both Python and R. The OTS data curation pipeline was written in Python and files were extracted and read using Pandas (70). Scikit-learn (71) was used to perform tSNE analyses of language model embeddings. Figure 1 was plotted in Python using the Matplotlib library (72). Figures 2-4 were plotted using ggplot and R tidyverse (73).

## Data Availability

The data within OTS is available under a CC-BY 4.0 license, freely downloadable and searchable at *https://opig.stats.ox.ac.uk/webapps/ots*. Datasets of public TCRs, chain coherence study clusters, and paired TCR language model weights are available from Zenodo *https://doi.org/10.5281/zenodo.11208211*.

## Code Availability

Python code for evaluating TCR chain coherence is available at *https://doi.org/10.5281/zenodo.11208211*. The paired TCR language model can be run by supplying the weights trained on TCRs to AbLang2 (*https://github.com/oxpig/AbLang2*).

## Author Contributions

MR, AGW, and CD conceptualized the study and designed the methodology. MR, AGW, and PA curated data. MR, AGW, BS, TO, NQ and OT developed software. MR, AGW, PA, and CD analysed the data. MR and AGW visualized the data. CD provided computational resources. MR and AGW wrote the original manuscript draft. CD reviewed/edited the manuscript and supervised the project.

## ACKNOWLEDGEMENTS

The work was supported through a Boehringer Ingelheim postdoctoral fellowship awarded to MR, research funding by Exscientia awarded to AGW, and Doctoral programme funding from the UK Engineering and Physical Sciences Research Council (EPSRC) awarded to TO, and OT (EP/S024093/1).

## Notes

https://zenodo.org/records/11208211

https://opig.stats.ox.ac.uk/webapps/ots

### Competing Interest Statement

The authors have declared no competing interest.

### Summary of Updates

Added a missing author. Nele P Quast. Her contribution was to software development.

