## Supplementary Figures and Tables for "The Observed T cell receptor Space database enables paired-chain repertoire mining, coherence analysis and language modelling"

### SI Figures and Tables

The following pages contain **4** SI Figures and **1** SI Tables.

*These authors contributed equally to this work

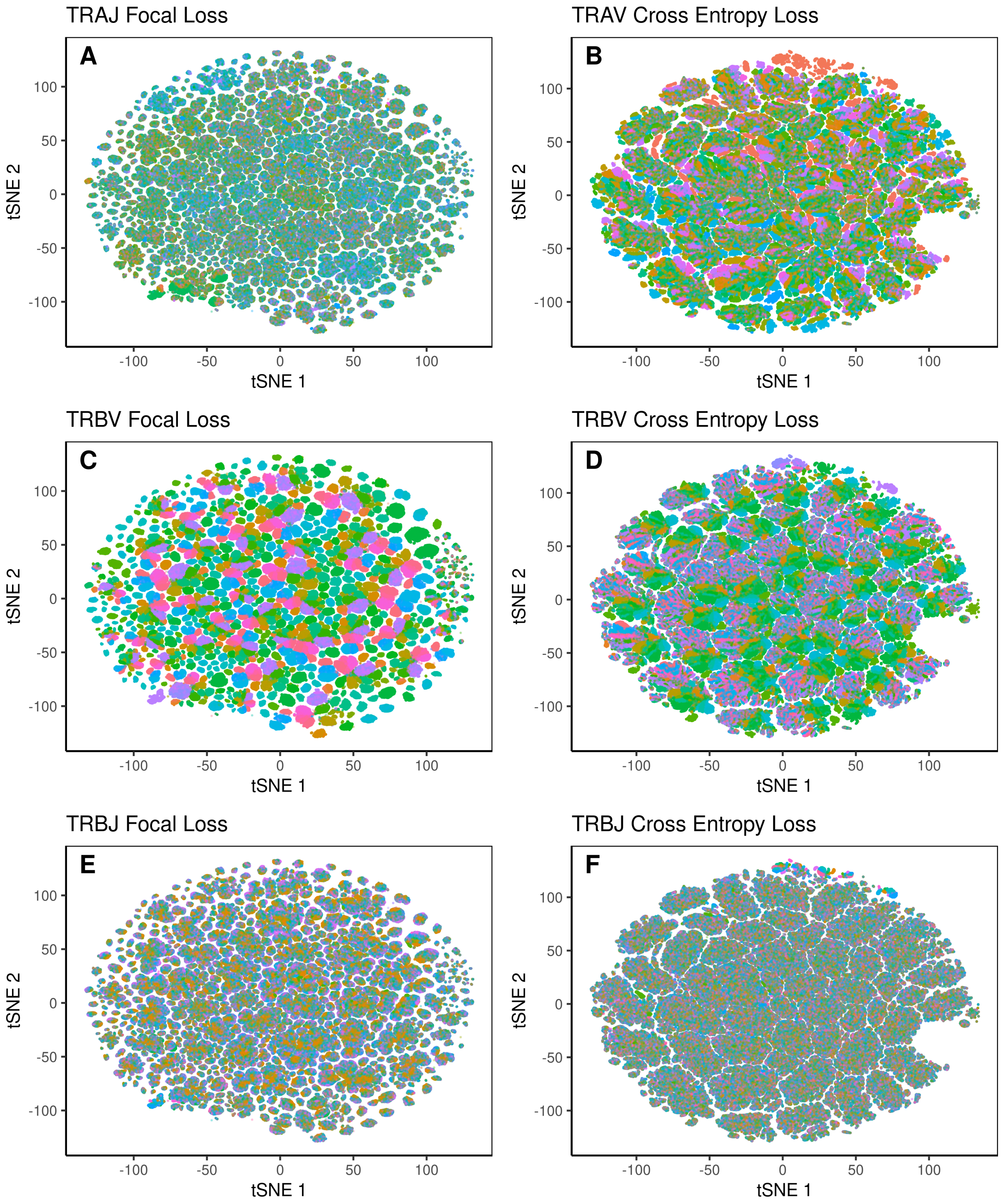


**SI Fig. 1. Focal loss and cross entropy loss embedding spaces annotated by VJ-gene usage.** Paired sequence embeddings were generated for 1.605M non redundant human TCR pairs in OTS using models trained with focal loss (A, C, E) or cross entropy loss (B, D, F) . The paired embedding space was visualised with tSNE analysis and points coloured by TRAJ (A), TRAV (B), TRBV (C-D) or TRBJ (E-F) gene usage.

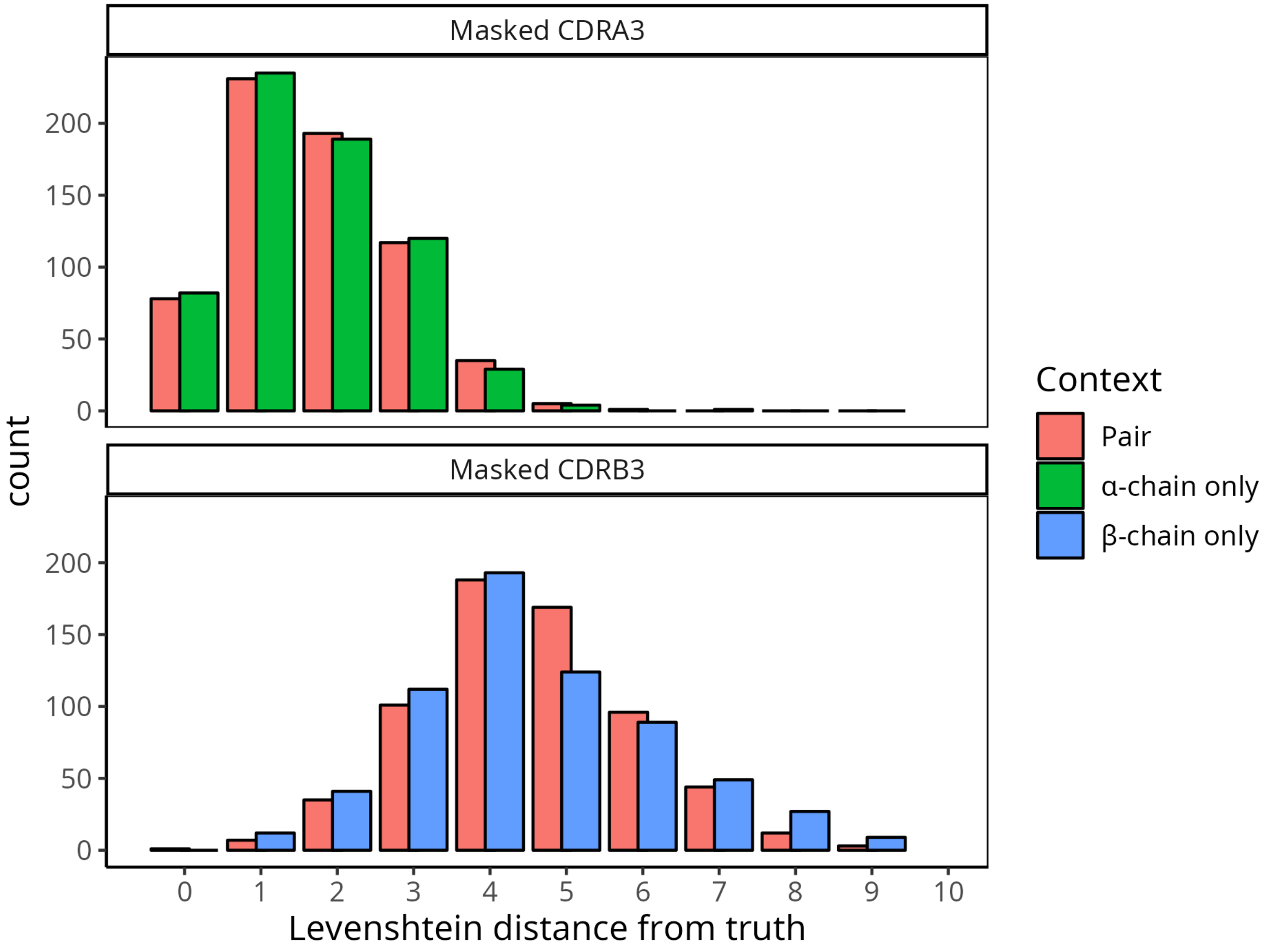


**SI Fig. 2. Sequence recovery distribution of masked MAIT TCR CDR3 sequences when given the context of only one, or both, chains.** MAIT TCRs not seen in paired OTS were collected from Garner et al. 2023 (1) and the CDR3 sequences fully masked. The distribution of Levenshtein distances of TCRLang-Paired infilled sequence from the ground truth is shown for three contexts. The paired context (both chains provided) is shown in red, $\alpha$-chain only is shown in green and $\beta$-chain only shown in blue. Top panel is from masking of the CDRA3, and bottom panel is the corresponding result for masking of the CDRB3.

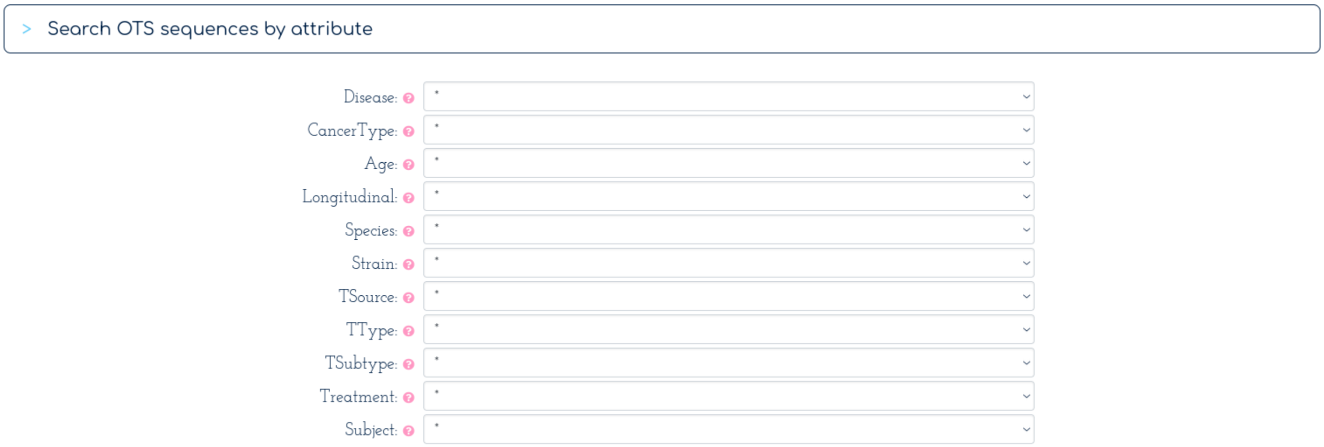


**SI Fig. 3. The OTS web app allows users to explore data in OTS.** Users can select studies matching fields of interest and subset based on a variety of parameters.

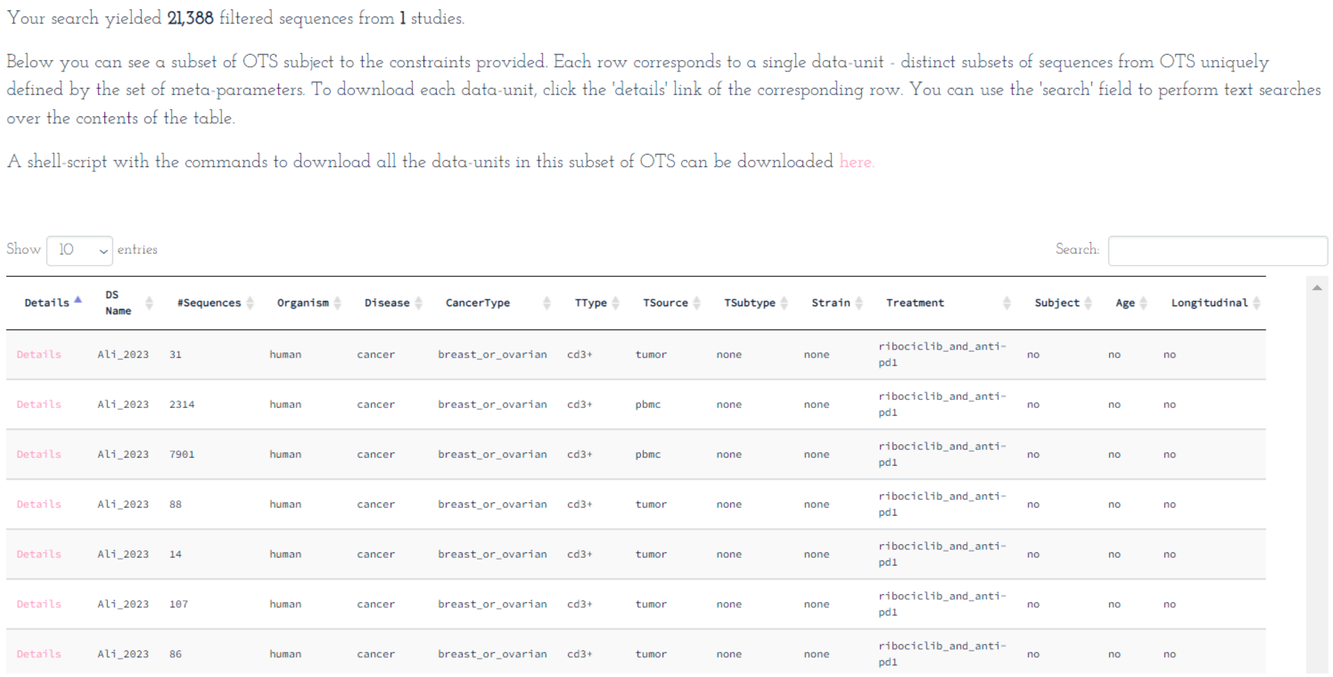
Your search yielded **21,388** filtered sequences from **1** studies.

Below you can see a subset of OTS subject to the constraints provided. Each row corresponds to a single data-unit - distinct subsets of sequences from OTS uniquely defined by the set of meta-parameters. To download each data-unit, click the 'details' link of the corresponding row. You can use the 'search' field to perform text searches over the contents of the table.

A shell-script with the commands to download all the data-units in this subset of OTS can be downloaded here.

Show 10 entries Search:

| Details | DS Name | #Sequences | Organism | Disease | CancerType | TType | TSource | TSubtype | Strain | Treatment | Subject | Age | Longitudinal |
|---|---|---|---|---|---|---|---|---|---|---|---|---|---|
| Details | Ali_2023 | 31 | human | cancer | breast_or_ovarian | cd3+ | tumor | none | none | ribociclib_and_anti-pd1 | no | no | no |
| Details | Ali_2023 | 2314 | human | cancer | breast_or_ovarian | cd3+ | pbmc | none | none | ribociclib_and_anti-pd1 | no | no | no |
| Details | Ali_2023 | 7901 | human | cancer | breast_or_ovarian | cd3+ | pbmc | none | none | ribociclib_and_anti-pd1 | no | no | no |
| Details | Ali_2023 | 88 | human | cancer | breast_or_ovarian | cd3+ | tumor | none | none | ribociclib_and_anti-pd1 | no | no | no |
| Details | Ali_2023 | 14 | human | cancer | breast_or_ovarian | cd3+ | tumor | none | none | ribociclib_and_anti-pd1 | no | no | no |
| Details | Ali_2023 | 107 | human | cancer | breast_or_ovarian | cd3+ | tumor | none | none | ribociclib_and_anti-pd1 | no | no | no |
| Details | Ali_2023 | 86 | human | cancer | breast_or_ovarian | cd3+ | tumor | none | none | ribociclib_and_anti-pd1 | no | no | no |

**SI Fig. 4. The OTS web app allows users to download relevant files for further analyses.** Search results are displayed for each file that fits the user defined search parameters. This figure shows the results of a search for "Disease: cancer, CancerType: breast_or_ovarian, Species: human". All or subsets of files that match the search criteria can then be downloaded.

| Redundant/Non-Redundant | CDR3 LD | Category | Mean Alpha Chain Coherence | Mean Beta Chain Coherence |
|---|---|---|---|---|
| Redundant | 0 | Baseline | **6.90%** (N=34142) | **4.11%** (N=166362) |
| | 1 | Baseline | 3.53% (N=19922) | 3.10% (N=48103) |
| | 2 | Baseline | 2.56% (N=9831) | 2.81% (N=15225) |
| | 0 | Tetramer | **44.33%** (N=124) | **42.97%** (N=887) |
| | 1 | Tetramer | 41.90% (N=110) | 25.89% (N=713) |
| | 2 | Tetramer | 40.24% (N=105) | 20.70% (N=595) |
| Non-redundant | 0 | Baseline | **6.49%** (N=34103) | **4.08%** (N=166310) |
| | 1 | Baseline | 3.44% (N=19885) | 3.10% (N=48082) |
| | 2 | Baseline | 2.54% (N=9792) | 2.81% (N=15209) |
| | 0 | Tetramer | **14.61%** (N=54) | **9.75%** (N=824) |
| | 1 | Tetramer | 14.00% (N=52) | 7.80% (N=666) |
| | 2 | Tetramer | 10.94% (N=50) | 6.26% (N=560) |

**SI Table 1.** Mean $\alpha$- and $\beta$-chain coherence values for the Baseline and Tetramer datasets from OTS. Redundant statistics are based on a dataframe that allows identical paired sequences to be found in different repertoires from the same study; these are filtered out to retrieve the non-redundant statistics. CDR3 Levenshtein Distance (LD) reflects the maximum number of edit distances between CDR3 sequences within a clonotype (2). Clonotypes and partner chain coherence are defined at the level of the allele.
